# Subtractive adaptation is a more effective and general mechanism in binocular rivalry than divisive adaptation

**DOI:** 10.1101/2023.01.19.524840

**Authors:** Maria Inês Cravo, Rui Bernardes, Miguel Castelo-Branco

## Abstract

The activity of neurons is influenced by random fluctuations and can be strongly modulated by firing rate adaptation, especially in sensory systems. Still, there is an ongoing debate about the characteristics of neuronal noise and the mechanisms of adaptation, and even less is known about how exactly they affect perception. Noise and adaptation play central roles in binocular rivalry, a visual phenomenon where two images compete for perceptual dominance. Here, we investigated the effects of different noise processes and adaptation mechanisms on visual perception by simulating a model of binocular rivalry with Gaussian white noise, Ornstein-Uhlenbeck noise, and pink noise, in variants with divisive adaptation, subtractive adaptation, and without adaptation. By simulating the nine versions of the model for a wide range of parameter values, we find that white noise only produces rivalry when paired with subtractive adaptation and that subtractive adaptation reduces the influence of noise intensity on rivalry strength and introduces convergence of the mean percept duration, an important metric of binocular rivalry, across all noise processes. In sum, our results show that white noise is an insufficient description of background activity in the brain and that subtractive adaptation is a stronger and more general switching mechanism in binocular rivalry than divisive adaptation, with important noise-filtering properties.

**Author Summary:** Visual neurons adapt to the environment by reducing the number of spikes evoked by a constant stimulus. They are also susceptible to random spikes produced by nearby neurons. These two phenomena, adaptation and noise, are essential features of brain activity and affect how we perceive the world. Although we know a great deal about the visual system, our understanding of the properties and mechanisms of neuronal noise and adaptation is still piecemeal, and even less is known about how these microscopic processes affect macroscopic behaviors. We shed light on this question by studying a bistable visual phenomenon called binocular rivalry, where two images compete for perception and where noise and adaptation play important roles. We simulated the activity of neurons involved in binocular rivalry to test different hypotheses about the statistics of neuronal noise and the mechanisms of adaptation. Our results reveal important differences between subtractive and divisive adaptation, suggesting that subtractive adaptation is a stronger switching mechanism in binocular rivalry and an effective noise filter. Our simulations also show the fundamental distinction between noise with and without temporal correlation, supporting the correlated noise hypothesis.

## 1. Introduction

Neuronal noise and firing rate adaptation are essential features of the neural activity that sustains brain function. Particularly, they affect the retrieval of information from sensory experience. However, the underlying mechanisms are not fully understood. Binocular rivalry is a visual phenomenon well poised to study noise and adaptation: when two images are presented simultaneously and independently to the two eyes, the neural representations of both stimuli compete for dominance, leading to an alternated perception of each image. This alternation has a stochastic structure and is proposed to occur due to adaptation to the dominant stimulus and random perturbations to neuronal activity (1).

Adaptation is a form of short-term plasticity induced by prolonged exposure to a stimulus. It manifests physiologically as a reduction in the spiking activity of sensory and cortical neurons and perceptually as altered sensitivity to certain stimulus features, such as reduced perceived contrast after adaptation to a high-contrast stimulus (2). Adaptation is beneficial for code efficiency because it allows the input-output response function of neurons to reflect the statistical properties of the environment by reducing sensitivity to constant stimuli and increasing responsiveness to less frequent stimuli (3). At the single-cell level, firing rate adaptation can result from hyperpolarizing potassium currents triggered by action potentials, which effectively raise the spiking threshold. At the network level, synaptic depression may reduce the drive to an adapting neuron. In contrast, changes in the non-classical receptive field, mediated by normalization to the activity of neighboring neurons, may modulate the gain function (4). The diversity of mechanisms and their distinct effects on the neuronal response function make it challenging to connect the perceptual effects of adaptation to their underlying neural representations.

At the single-neuron level, adaptation mechanisms can have different temporal profiles and computational effects. Experimental recordings of simple and complex cells in the primary visual cortex (V1) of cats (5) and of neurons in the middle temporal visual area (MT) of monkeys (6) have documented an exponential decay of the firing rate during adaptation. Other studies, including those of the fly visual system (7), the somatosensory barrel cortex of mice (8) and electrosensory neurons of electric fish (9), have reported that the firing rate has a power-law dependence on time. This power-law behavior may be due to the existence of multiple exponential adaptation processes with different time scales (2,10). Indeed, several mechanisms activate the somatic hyperpolarization currents that follow an action potential, including sodium- and calcium-gated potassium currents (4,11,12). Simulation studies have shown that these currents affect the neuronal response function differently: calcium-gated currents have a divisive effect on the input-output function, while sodium-gated currents have a predominantly subtractive effect on the response function, shifting it laterally to higher input values and functioning as a high-pass filter (13). The subtractive and divisive effects of adaptation allow the neuron to adapt to the mean and the variance of the incoming signal, respectively (14). Taken together, these findings have led computational models of visual perception to include adaptation as an exponential process that depends linearly on the firing rate and that either subtracts from the synaptic input (subtractive adaptation) or adds to the denominator of the nonlinear input-output response function (divisive adaptation).

Neuronal noise consists of random perturbations to the spiking activity of neurons, mainly due to fluctuations in the release, diffusion and binding of neurotransmitters in the synapse. There are also fluctuations in the membrane potential (thermal noise) and in the opening and closing of ion channels (electrical noise) (15). Still, synaptic noise is the most significant contributor to variability in neural activity (16,17). By providing the background activity against which signals are superimposed, synaptic noise affects information transmission, most commonly through deterioration of the signal-to-noise ratio, although in some cases it can improve the detectability of a well-matched signal through stochastic resonance (18,19). Since it is present throughout the visual system, noise can cause variability in perception when an observer is presented with the same stimulus twice (20). However, the impact of the statistical properties of neuronal noise on visual perception is still poorly understood.

The temporal structure of neuronal noise is its main statistical property that can be quantified in experimental studies, since intricate synaptic connectivity makes it difficult to separate signal from noise to quantify the mean level of synaptic background activity. Recordings of spike trains in area MT of monkeys showed that the frequency power spectrum was relatively flat and consistent with a Poisson process with a refractory period (21). In contrast, recordings in cat V1 and monkey inferior temporal (IT) area showed greater power at lower frequencies and a monotonic decrease at higher frequencies (22). Measurements of collective neural activity in the human brain, using techniques such as electroencephalography (EEG), magnetoencephalography (MEG), local field potential recordings and the blood-oxygen-level-dependent (BOLD) signal in resting-state functional magnetic resonance imaging (fMRI), also show non-Poisson power spectra scaling between 1/*f* and 1/*f*^2^(23–25). The statistics of spontaneous activity are at the heart of the debate on whether information is encoded by the precise timing of spikes or the average firing activity of neurons. Poisson statistics are associated with a rate code, while non-Poisson statistics, with a temporal correlation between successive spikes, are associated with a temporal code. Neurons may operate in both regimes (26,27). Furthermore, temporally uncorrelated spike trains can give rise to temporally correlated activity measures through the nonlinear filtering properties of both dendrites (28,29) and extracellular space (30). As a result, computational models have included neuronal noise as a synaptic current of zero-mean Gaussian white noise (an approximation to a Poisson process), pink or Brownian noise (power spectra weighted as 1/*f* and 1/*f*^2^, respectively), and Ornstein-Uhlenbeck noise (a low-pass filtered version of white noise, which is related to Brownian noise).

Computational models of binocular rivalry have included different implementations of neuronal noise and firing rate adaptation. Noise has been modeled as an Ornstein-Uhlenbeck process (31–34), a pink noise process (35) and a white noise process (36–39), either as a stochastic synaptic current or as perturbations to auxiliary variables like adaptation (37,40) and synaptic depression (41). Adaptation is often modeled as an exponential process in subtractive (34,39) or divisive (42,43) formulations. Two studies have made comparisons of binocular rivalry models with different descriptions of noise and adaptation: Baker and Richard (35) compared white, pink, and Brownian noise and cases in between, using subtractive adaptation, while Shpiro et al. (44) compared subtractive and divisive adaptation with synaptic depression using Ornstein-Uhlenbeck noise.

Here, we study the influence of different noise and adaptation mechanisms on visual perception by simulating a computational model of binocular rivalry. We compare Gaussian white noise, pink noise and Ornstein-Uhlenbeck noise in versions of the model with (divisive or subtractive) and without adaptation. Simulation of the model for a range of noise amplitudes and visual contrasts revealed that white noise only gives rise to strong binocular rivalry when paired with subtractive adaptation. In contrast, correlated noise generates binocular rivalry dynamics in a larger region of parameter space, regardless of adaptation. Simulations with subtractive adaptation also show a reduced influence of noise intensity and similar values of mean percept duration across noise processes. Comparison of simulated dynamics with known constraints on mean percept duration enabled the determination of the minimum temporal correlation of Ornstein-Uhlenbeck noise.

## 2. Results

We study the role of noise statistics and adaptation mechanisms on visual perception by simulating a binocular rivalry model with monocular, binocular and ocular opponency neurons (Fig. 1A). We include additive synaptic noise as an independent stochastic source for each neuron and compare three stochastic processes: Gaussian white noise, Ornstein-Uhlenbeck noise, and pink noise (Fig. 1B). We include firing rate adaptation as either subtractive feedback to the synaptic current entering the soma (subtractive adaptation) or as an increase in the saturation of the nonlinear input-output function that transforms synaptic current into firing rate (divisive adaptation), and we compare these formulations with the model without adaptation. Simulations were run for a range of parameter values to explore parameter space and detect phase transitions. We changed the contrast of the input images to the two eyes (*c*), the intensity of the noise process (*σ*), and, for Ornstein-Uhlenbeck noise, the correlation time (*τ*). For each simulation we calculated the relative dominance time (RDT), a measure of rivalry strength, the mean dominant percept duration, the coefficient of variation of percept durations, and the mean mixed percept duration (see Methods). These quantities were averaged over three runs of 60 s for each pair of parameters, and the respective standard deviation was calculated.

**Fig. 1:**
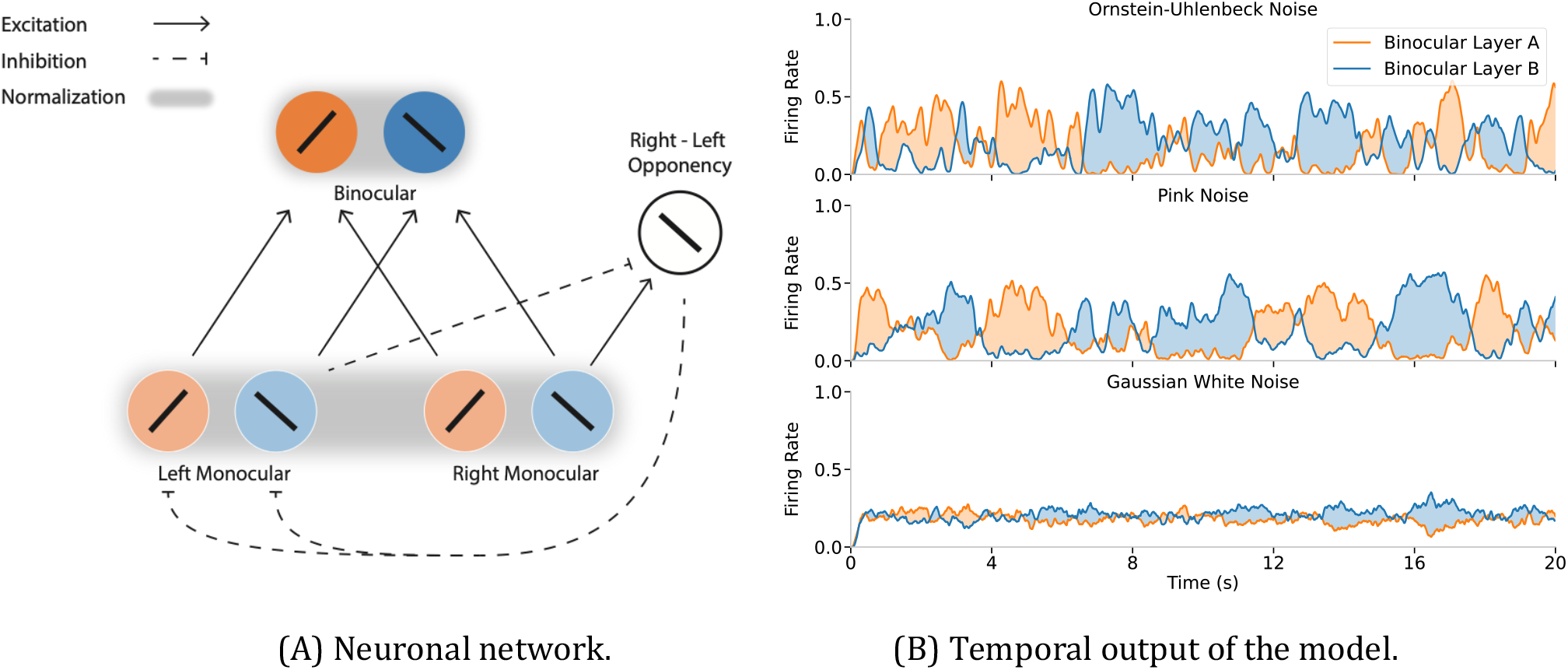
Model of binocular rivalry used in the simulations. (A) Network architecture with three layers: monocular neurons, binocular neurons, and opponency neurons (33). Only one of the four types of opponency neurons is shown. (B) Simulated activity of binocular neurons responsive to different patterns for three models of internal noise.

### White noise only leads to strong rivalry with subtractive adaptation

We first investigate if the model predicts similar rivalry strengths for each type of noise and how adaptation affects these results (Fig. 2). For a model without adaptation, there is a large region of parameter space with high dominance of one percept over the other, but only if synaptic noise is an Ornstein-Uhlenbeck or a pink noise process. In these cases, the dominance increases with noise intensity, especially for intermediate input contrast. For white noise, no pair of parameters leads to dominance stronger than 36%.

**Fig. 2:**
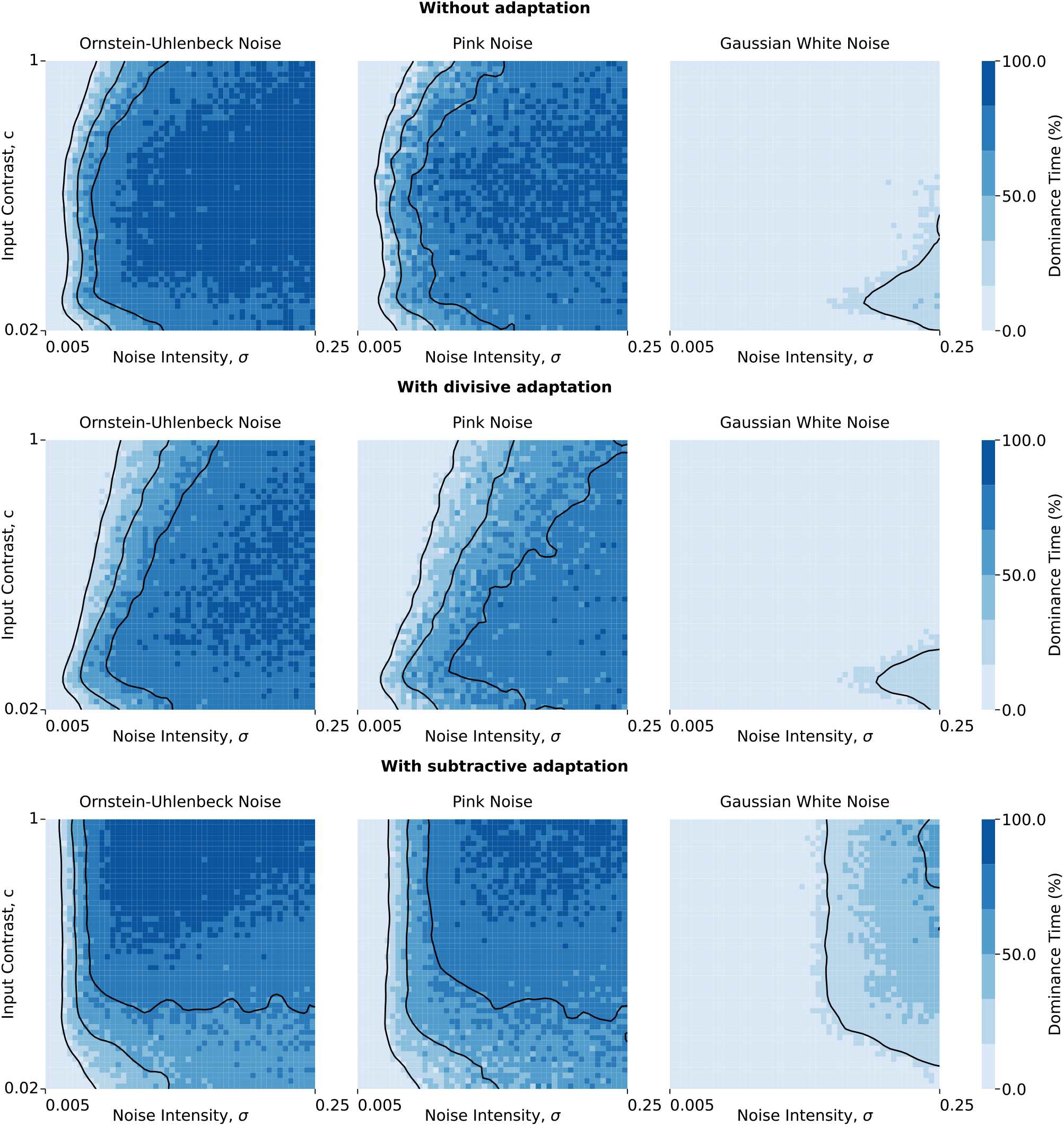
Effect of noise intensity and input contrast on relative dominance time, for models with and without adaptation. Darker squares denote strong dominance and lighter squares denote mixed perception. Each square is the average of three simulations. Contours correspond to RDT equal to 20%, 50% and 70% and were smoothed with a Gaussian filter. For simulations with Ornstein-Uhlenbeck noise, the temporal correlation of noise was kept constant at *τ* = 500 ms. For simulations with adaptation, adaptation weight and adaptation time scale were kept constant at *w*_*H*_ = 2 and *τ*_*H*_ = 2000 ms, respectively.

When the model includes divisive adaptation, the dominant region in the diagrams for temporally correlated noise shrinks and shifts to the right towards higher noise intensity values. The diagram for uncorrelated noise remains largely unchanged.

When the model includes adaptation as a subtractive synaptic input, the dominant region for correlated noise shifts upwards and rivalry strength is shaped more by input contrast than by noise intensity. Interestingly, this type of adaptation gives rise to stronger rivalry for uncorrelated noise. Taken together, these simulations show that with white noise only subtractive adaptation leads to strong rivalry. This finding is robust to changes in adaptation parameters and in the mathematical formulation of divisive adaptation (see Supplementary Figures).

### Subtractive adaptation softens the difference between noise processes for mean percept duration

We find the results depicted in Fig. 3 when examining the effect of adaptation on other metrics of rivalry and considering only cases with sufficiently strong rivalry, defined by relative dominance time above 50%. The statistical significance of pairwise comparisons was assessed with Mann-Whitney U tests with a Bonferroni correction.

**Fig. 3:**
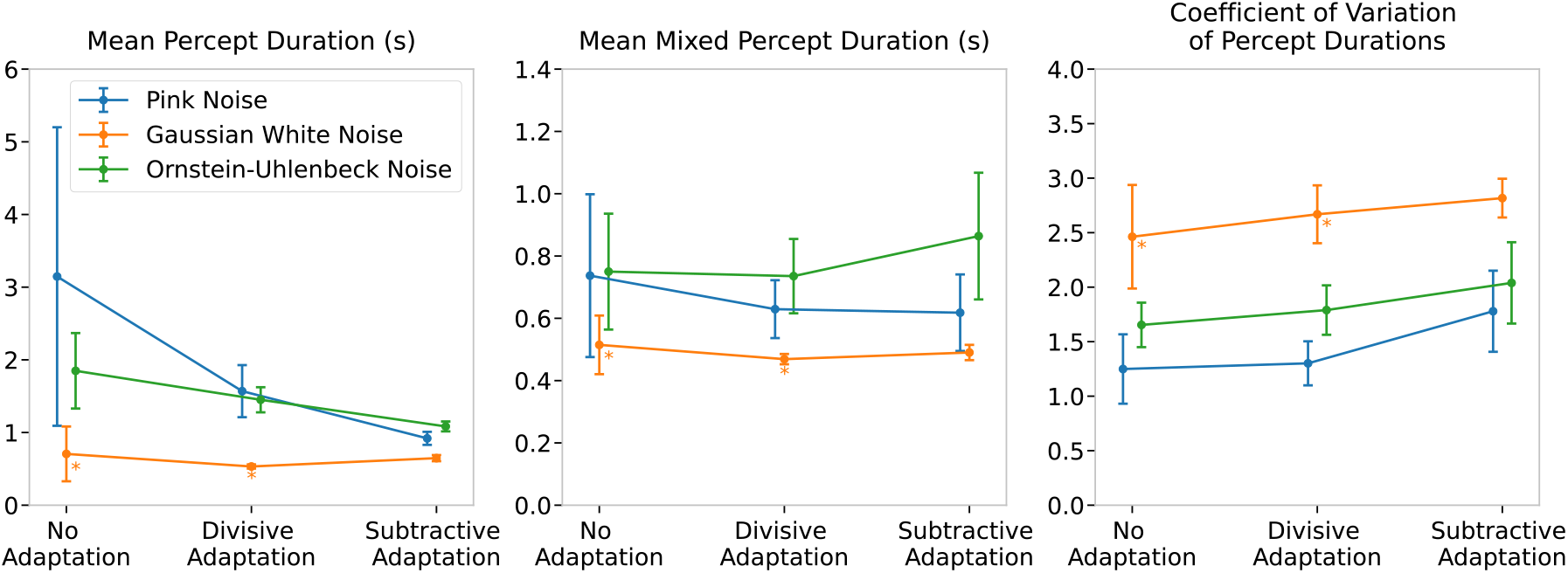
Effect of adaptation on binocular rivalry metrics for each noise process. Mean and standard deviation of three binocular rivalry metrics. Only cases with RDT > 50% and for which each metric is defined (see Methods) are included. For white noise without adaptation and with divisive adaptation (marked with an asterisk), there are no cases with RDT > 50%, so this constraint was lifted and RDT > 20% was used instead (see Table S1).

For pink and Ornstein-Uhlenbeck noise processes, adaptation reduces the mean duration of dominant percepts (*p* < 0.001 for both adaptation formulations), as is physiologically expected. Subtractive adaptation leads to a stronger reduction (*p* < 0.001) and results in similar values of this metric for all noise processes.

The effect of adaptation on mean mixed percept duration is not consistent. Mixed perception increases significantly for Ornstein-Uhlenbeck noise when subtractive (*p* < 0.001), but not divisive (*p* > 0.5), adaptation is added, whereas for pink noise adaptation reduces mixed perception (*p* < 0.001 for both adaptation variants). The coefficient of variation of percept durations increases with divisive and subtractive adaptation for all noise processes (*p* < 0.05 for all comparisons).

### Correlated noise generates stronger rivalry and longer and more reliable dominance periods than uncorrelated noise

The cumulative histogram of relative dominance time for simulations with subtractive adaptation (Fig. 4A) highlights the contrast between correlated and uncorrelated noise processes. Although the distributions of both types of correlated noise closely follow each other, Ornstein-Uhlenbeck noise displays stronger dominance. The plot in Fig. 4B shows the difference in binocular rivalry metrics between the three noise processes for the region of parameter space where relative dominance time is higher than 50%. Percept durations are longer for Ornstein-Uhlenbeck noise and shorter for white noise. The same pattern of results occurs in mean mixed percept duration, although with larger differences in the variation of this metric. White noise generates dominant percepts with more varied durations, as seen by the coefficient of variation.

**Fig. 4:**
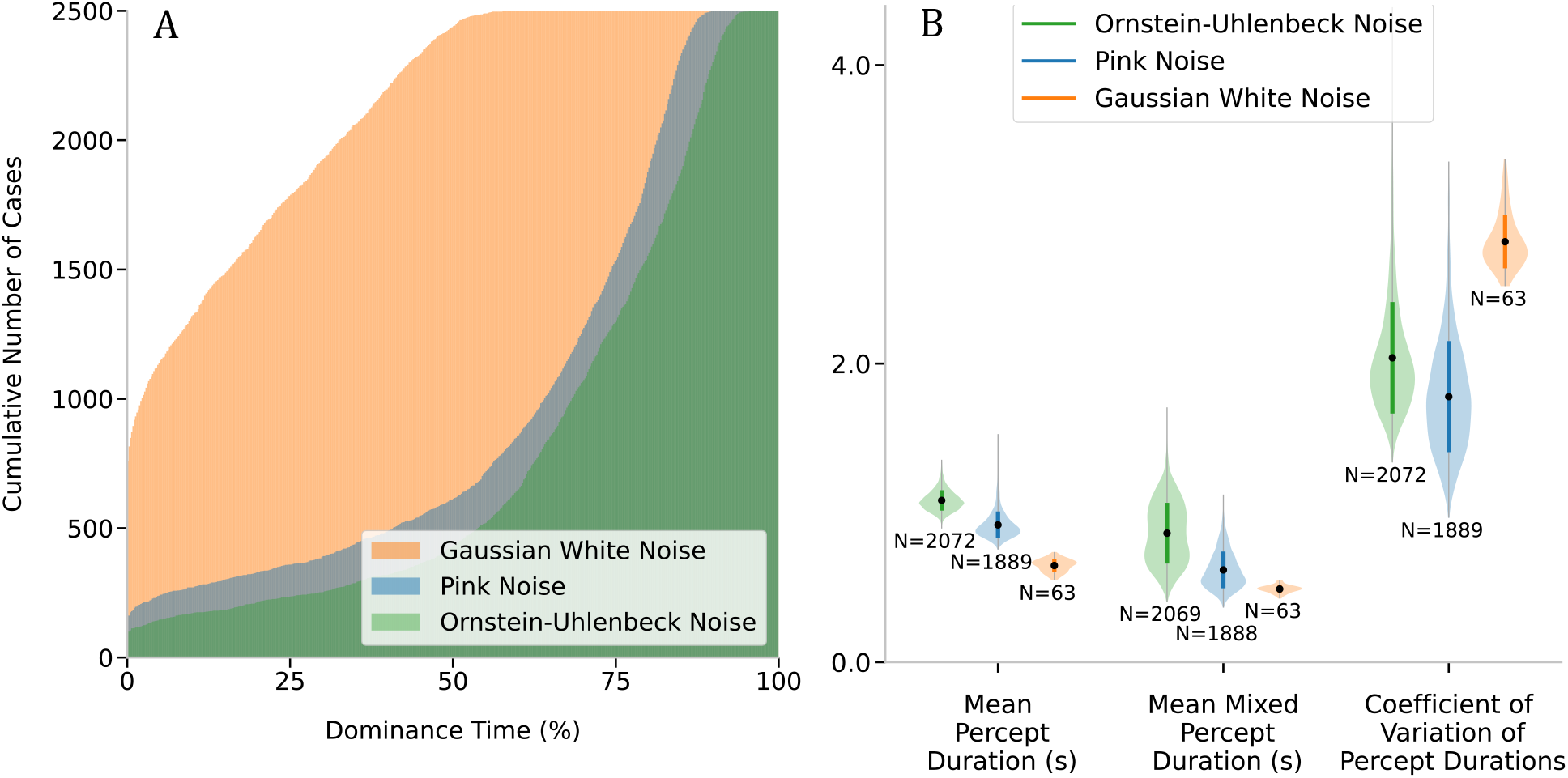
Distributions of rivalry metrics for simulations with subtractive adaptation. (A) Cumulative histogram of relative dominance time of three types of noise, using the data of Fig.2. (B) Mean, standard deviation and density kernel estimation of mean duration of dominant and mixed percepts, in seconds, and the coefficient of variation of dominant percept durations for simulations with Ornstein-Uhlenbeck noise, pink noise and Gaussian white noise, calculated over the ranges of image contrast and noise intensity that satisfy *RDT* > 50% and for which the metrics are defined, resulting in different N values (see Methods and Table S1).

### The minimum correlation time for Ornstein-Uhlenbeck noise is 400 ms when combined with subtractive adaptation

Ornstein-Uhlenbeck noise is defined by two parameters, noise intensity (*σ*) and correlation time (*τ*), so we simulated the model for a range of values of *σ* and *τ* (Fig. 5). For simulations without adaptation, the relative dominance time increases with noise intensity and noise correlation time, whereas in simulations with subtractive adaptation noise intensity is the determining factor. By superimposing the region of parameter space where mean dominant percept duration is above the experimental minimum of 1 s (45–47), we can determine the minimum correlation time for Ornstein-Uhlenbeck noise. Without adaptation, the mean dominant percept duration is above 1 s for all pairs of parameters tested. However, when adaptation is introduced, the mean percept duration is above 1 s for *τ* ≳ 400 ms. The right panels of Fig. 5 show that the standard deviation of the relative dominance time diverges in a specific parameter space region, defining a border between weak and strong rivalry. Considering the relative dominance time as the order parameter of this dynamical system, we conclude that there is a phase transition governed by *σ* and *τ*, with *σ* dominating, especially for the model with subtractive adaptation.

**Fig. 5:**
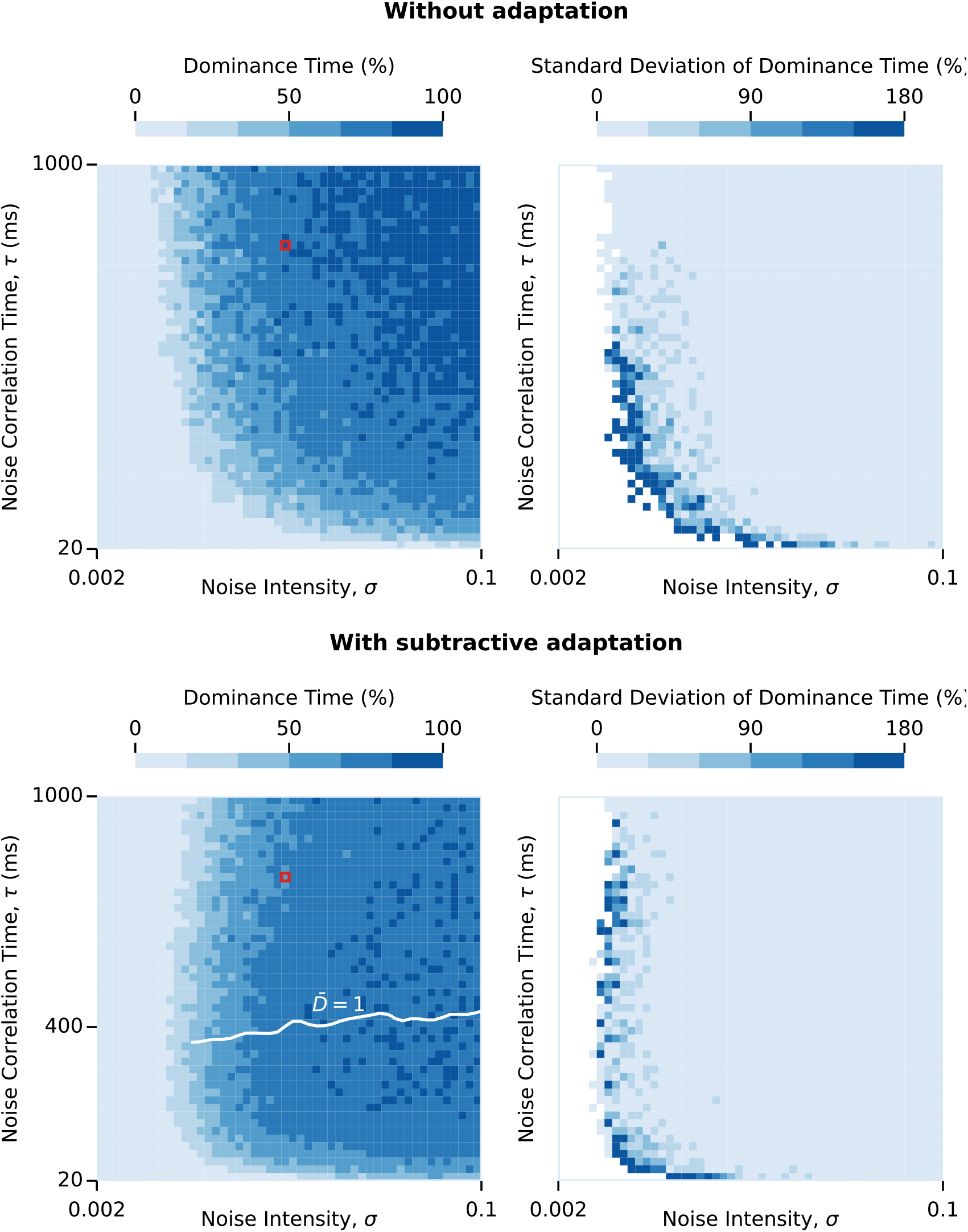
Ornstein-Uhlenbeck noise parameter space. Effect of noise intensity *σ* and noise correlation time *τ* on relative dominance time, *RDT*, and percent standard deviation of relative dominance time, *δRDT*/*RDT*, over three simulations. The red squares mark the parameters used by the original authors of the model (33). The white line marks the boundary above which the mean dominant percept duration is larger than 1 s. For simulations with adaptation, adaptation weight and adaptation time scale were kept constant at *w*_*H*_ = 2 and *τ*_*H*_ = 2000 ms, respectively.

## 3. Discussion

Simulations of binocular rivalry with three types of neuronal noise and two types of adaptation showed remarkable differences between correlated and uncorrelated synaptic noise and between subtractive and divisive adaptation. Subtractive adaptation is the only condition under which white noise can give rise to strong binocular rivalry. It also changes the dynamics with correlated noise such that noise intensity loses impact compared to simulations without and with divisive adaptation. By reducing the mean percept duration predicted with Ornstein-Uhlenbeck noise and pink noise, subtractive adaptation converges the value of this metric for all noise processes. When comparing simulations with subtractive adaptation, Ornstein-Uhlenbeck noise generates stronger rivalry and longer dominant and mixed percept durations, while Gaussian white noise generates more variability in dominant percepts. Finally, comparing simulations with Ornstein-Uhlenbeck noise and subtractive adaptation with experimental constraints on the minimum percept duration allowed us to determine the minimum correlation time for Ornstein-Uhlenbeck noise.

Our computational study reveals profound differences between divisive and subtractive adaptation. First, the results for Gaussian white noise show that subtractive adaptation is a much stronger switching mechanism than divisive adaptation. Second, while neuronal noise is still necessary for rivalry, subtractive adaptation shifts the system away from noise-driven behavior, which is in line with the conclusions of Shpiro et al. (34) that binocular rivalry requires a balance between adaptation and noise. Third, by blurring the difference between mean percept duration for all three noise processes, subtractive adaptation functionally filters neuronal noise, reducing its impact on visual perception. Together, these results suggest that subtractive adaptation is a more effective and general mechanism in binocular rivalry than divisive adaptation.

The functional differences between subtractive and divisive adaptation are explained by their effects on the neuronal input-output function. Subtractive adaptation shifts the response function towards higher input values, rendering the neuron entirely insensitive to small fluctuations. Meanwhile, divisive adaptation scales down the response function, introducing a small change in responsiveness. The relative effect of divisive adaptation is further reduced when the pool of normalization neurons is active, which happens continually in this model of binocular rivalry or indeed in any model of a visual phenomenon where there is competition through divisive inhibition between the neuronal representations of different stimuli. It is, therefore, reasonable to expect that our findings on the effectiveness and noise-filtering properties of subtractive adaptation generalize to other models of visual function. Particularly, our results suggest that subtractive adaptation could have a determinant role, perhaps as the dominant adaptation mechanism, in areas of the nervous system where denoising is more important than transmitting precise temporal information and in areas where flexible neural representations are advantageous.

Although subtractive adaptation harmonizes our results for different types of noise, there are still distinctions, particularly between correlated and uncorrelated noise processes. Simulations with pink noise and Ornstein-Uhlenbeck noise give rise to longer and less variable dominance periods, although the coefficient of variation is still far from the experimental range, between 0.4 and 0.6 (45,46). Furthermore, these stochastic processes generate binocular rivalry for weaker noise intensities, while Gaussian white noise requires noise strengths close to 0.25. This means that if internal noise in the brain is temporally uncorrelated, it would have to operate at higher levels to influence perception. Our results thus support the correlated noise hypothesis, but unlike Baker and Richard (35), who concluded that pink noise was a better model, we are unable to make a clear distinction between the two. However, our work provides some constraints: besides virtually excluding white noise, it establishes a minimum temporal correlation constant for Ornstein-Uhlenbeck noise, which should inform future computational studies on neural activity.

In conclusion, by simulating a binocular rivalry model with different noise processes and adaptation mechanisms for a wide range of parameter values, we showed that subtractive adaptation is a better candidate for a switching mechanism in binocular rivalry and that correlated noise is a better candidate for the distribution of spontaneous activity in the brain. Our work contributes to the understanding of firing rate adaptation by demonstrating the noise-filtering properties of subtractive adaptation at the level of perception.

## 4. Methods

To simulate binocular rivalry, we used a model proposed by Said and Heeger (33), which relies on ocular opponency neurons to detect interocular conflict. The dynamics are described by a two-step firing rate model, with one differential equation describing the synaptic current delivered to the soma of the neurons and another equation for the postsynaptic firing rate as a function of this current. The architecture of the network determines the inputs to each neuron. Because this is a firing rate model, each unit in the network can represent a population of neurons, and the computed firing rate can be interpreted as the mean firing rate of a population of similar neurons (48). The original model only relied on neuronal noise as the sole switching mechanism, and here we added firing rate adaptation as an extra differential equation for each unit.

### Neuronal Network Model

The model considers two orthogonal orientations, which constitute the competing stimuli. The model has three types of neurons: monocular, binocular and ocular opponency neurons (Fig. 1A). Left and right monocular neurons receive visual input from each eye, depending on orientation preference. Binocular neurons sum the activity from the right and left monocular neurons with the same orientation preference. Opponency neurons subtract the activity of left and right monocular neurons with the same orientation preference such that there are two right-minus-left (R-L) opponency neurons, one for each orientation, and two left-minus-right (L-R) neurons. The opponency neurons then inhibit all monocular neurons from the side which was subtracted, i.e., R-L opponency neurons inhibit left monocular neurons, including the population with orthogonal orientation preference. The activity of binocular neurons is a proxy for perception (Fig. 1B).

There is no direct inhibition between monocular or binocular neurons. Instead, the inhibition within each neuronal layer occurs through divisive normalization of the synaptic input (see Eq. 3 and details below). There are four normalization pools: one for binocular neurons, composed of both binocular neurons, one for monocular neurons, composed of all four monocular units, and two for opponency neurons, the pool for R-L neurons and the one for L-R neurons. Furthermore, there is direct inhibition through the feedback from opponency neurons to the monocular neurons that were subtracted (dashed arrow in Fig. 1A).

### Differential Equations

The two-step firing rate model proposed by Said and Heeger (33) describes the exponential filtering of the presynaptic drive and defines the firing rate as a low-pass filtered version of the synaptic current. The synaptic drive is exponentially filtered by the dynamics of the synaptic conductance in response to a presynaptic spike and by the passive and active properties of the dendritic cables that carry the synaptic current to the soma of the neuron (48). The equation for the synaptic current *I*is:

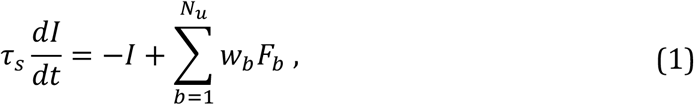

where *F*_*b*_ are the firing rates of the *N*_*u*_ presynaptic input neurons, with *w*_*b*_ the weights of these inputs and *τ*_*s*_ the time constant of the synapse-to-soma process. The architecture of the network determines the input sum. Monocular neurons receive sensory input which is added to the right-hand side (RHS) of the equation as input contrast.

The effects of membrane capacitance and resistance on membrane potential lead to low-pass filtering of the synaptic current (48). The postsynaptic firing rate *F* is thus described by:

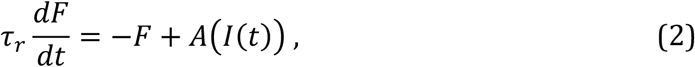

with *A(I)* an activation function, usually nonlinear, that describes the input-output function of the neuron and *τ*_*r*_ the time constant of this process, determining how closely *F* can follow fluctuations in *I*.

The activation function used in this model of binocular rivalry describes divisive input normalization (49):

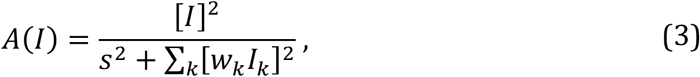

where [.] denotes half-wave rectification, *s* is a semi-saturation constant, which can be different for each type of neuron, and the weighted sum over *k* is the sum of the synaptic currents of the neurons in the normalization pool.

If the synaptic time constant *τ*_*s*_and the firing rate time constant *τ*_*r*_are significantly different, the system formed by Eqs. 1 and 2 can be replaced by only one differential equation (48). For instance, if *τ*_*r*_ ≪ *τ*_s_, the firing rate *F* follows *I*almost instantaneously and *F*(*t*) = *A*< *I*(*t*)), leaving only the differential equation for the synaptic current *I*. If *τ*_*r*_ ≫ *τ*_s_, the synaptic current reaches equilibrium faster than the firing rate and one can make the replacement 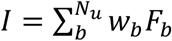, working only with the equation for the firing rate *F*. This last simplification is common and done in models of binocular rivalry by Shpiro et al. (34), Li et al (43) and Wilson (42), who use time constants *τ*_*r*_ of 10 ms or 20 ms, therefore assuming *τ*_*s*_ ≪ 10 ms. In contrast, Said and Heeger (33) use both equations with equal time constants *τ*_*s*_ = *τ*_*r*_ = 50 ms. This is also our approach.

### Noise

Noise is introduced in the equations by adding a stochastic process as a synaptic input on the RHS of Eq. 1. Each neuron has an independent noise source. This stochastic process can be an Ornstein-Uhlenbeck process, a Gaussian white noise process or a pink noise process.

The Ornstein-Uhlenbeck noise process η(*t*) is defined by a differential equation of a low-pass filter of a white noise process, ξ(*t*):

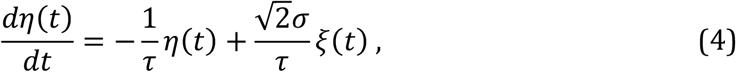

with *τ* the correlation time and *σ* the standard deviation of the Gaussian white noise process ξ(*t*). Ornstein-Uhlenbeck noise is thus exponentially-filtered white noise and models the low-pass filtering effects of synapses (34). However, instead of the synaptic time constant *τ*_*s*_ above, a much larger value for *τ* is usually chosen: *τ* = 800 ms in Said and Heeger (33) and *τ* = 100 ms in Shpiro et al. (34) and Li et al. (43). Integrating Eq. 4 along with the system equations significantly slows down the simulation. Hence, an alternative way of computing the Ornstein-Uhlenbeck process is by starting with Gaussian white noise of standard deviation *σ*, computed for all time steps, and convolving in time with a Gaussian kernel with standard deviation *τ*, as done by Said and Heeger (33). Although an exponential kernel would be more consistent with Eq. 1, a comparison of simulations with each type of kernel showed no significant differences.

For simulations with white noise instead of Ornstein-Uhlenbeck noise, the perturbations are sampled from a Gaussian distribution with standard deviation *σ*. Pink noise is computed by applying an inverse Fourier transform to a random process created in the frequency domain with amplitude proportional to 1/*f* and phase sampled from a uniform distribution, *φ* ∈ (0, 2*π*). To obtain the specified standard deviation in the time domain, the resulting process is multiplied by a correcting factor.

### Adaptation

We modelled adaptation as a slow exponential process that was either subtracted from the synaptic input of each neuron (subtractive adaptation) or added to the denominator of the activation function (divisive adaptation). The exponential dynamics of the adaptation variable *H* obey

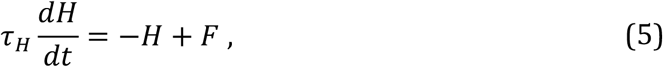

where *τ*_*H*_ is the adaptation time constant, and *F* is the neuron’s firing rate. The different formulations used are summarized in Table 1.

**Table 1:** Different mathematical implementations of firing rate adaptation used in simulations.

| Formulation | Added to RHS of<br>Eq.1 | Added to<br>denominator of<br>Eq.3 | Multiplied by<br>F in Eq.5 | Simulation |
| --- | --- | --- | --- | --- |
| 1 | 0 | $H$ | $w_H$ | Figs. 2, 3, S1, S2 |
| 2 | $-w_H H$ | 0 | 1 | Figs. 2-5, S1 |
| 3 | 0 | $w_H H$ | 1 | Fig. S1 |
| 4 | 0 | $w_H H^2$ | 1 | Fig. S1 |
| 5 | 0 | $H^2$ | $w_H$ | Fig. S1 |
| 6 | 0 | $w_H H^2$ | 1 | Fig. S1 |

Unless otherwise stated, we set adaptation parameters to *w*_*H*_ =2and *τ*_*H*_ = 2000 ms. These are the same values used by Li et al. (43). Shpiro et al. (34) also use *τ*_*H*_ = 2000 ms, while Wilson (42) uses *τ*_*H*_ = 900 ms.

### Metrics of Rivalry

In each simulation, binocular rivalry metrics are calculated based on the firing rate of binocular neurons, which is a proxy for the perception of one pattern or the other. Percepts were classified as dominant if they lasted longer than 300 ms and had a dominance index above 0.3 (31,43). The dominance index is defined as 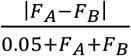. Using this classification, periods longer than 300 ms but with a dominance index below 0.3 are considered mixed percept periods. These periods are then used to calculate the mean dominant percept duration, the mean mixed percept duration, and the coefficient of variation of dominant percept durations, which is the ratio of the standard deviation of dominant percept durations to the mean dominant percept duration. Relative dominance time (RDT) is the fraction of simulated time where one percept is dominant.

The minimum duration of 300 ms is used by Moreno-Bote et al. (31) and Li et al. (43) and is based on the assumption that stimuli shorter than 300 ms are not perceived. The dominance index is used by Said and Heeger (33) and Li et al. (43) and mimics the definition of Michelson contrast (50) but we added the constant 0.05 to the denominator to avoid large values of the index when the firing rates of both neuronal populations are very close to zero, which happens when input contrast is low. This addition can be thought of as a minimum firing rate for visual perception and only significantly affects the measured strength of binocular rivalry for low contrasts.

### Simulation Details

The network contains ten neuronal units, which resulted in 20 differential equations in the model without adaptation and 30 when adaptation was included. These were integrated using MATLAB’s ODE45 routine (version R2020a), with noise introduced as an external function which was interpolated by the integration scheme. This interpolation was a good approximation of the stochastic process for all conditions tested (low contrast, high noise intensity, uncorrelated noise, etc.). Examples of simulated time series of binocular rivalry are presented in Fig. 1B.

By varying noise intensity *σ*, input contrast *c* and correlation time *τ* (in the case of Ornstein-Uhlenbeck noise), the dynamics of binocular rivalry were simulated for 2500 pairs of parameters in each diagram (Table 2), with metrics of rivalry calculated for each simulation of 60 seconds and averaged over three runs. The corresponding standard deviation over the three runs was calculated. The duration of each simulation and the number of repetitions were chosen to obtain enough variability without significantly slowing the simulations since 7500 simulations were performed to obtain each diagram. The list of parameter values kept constant and equal to the original values used by Said and Heeger (33) is shown in Table 3.

**Table 2:**
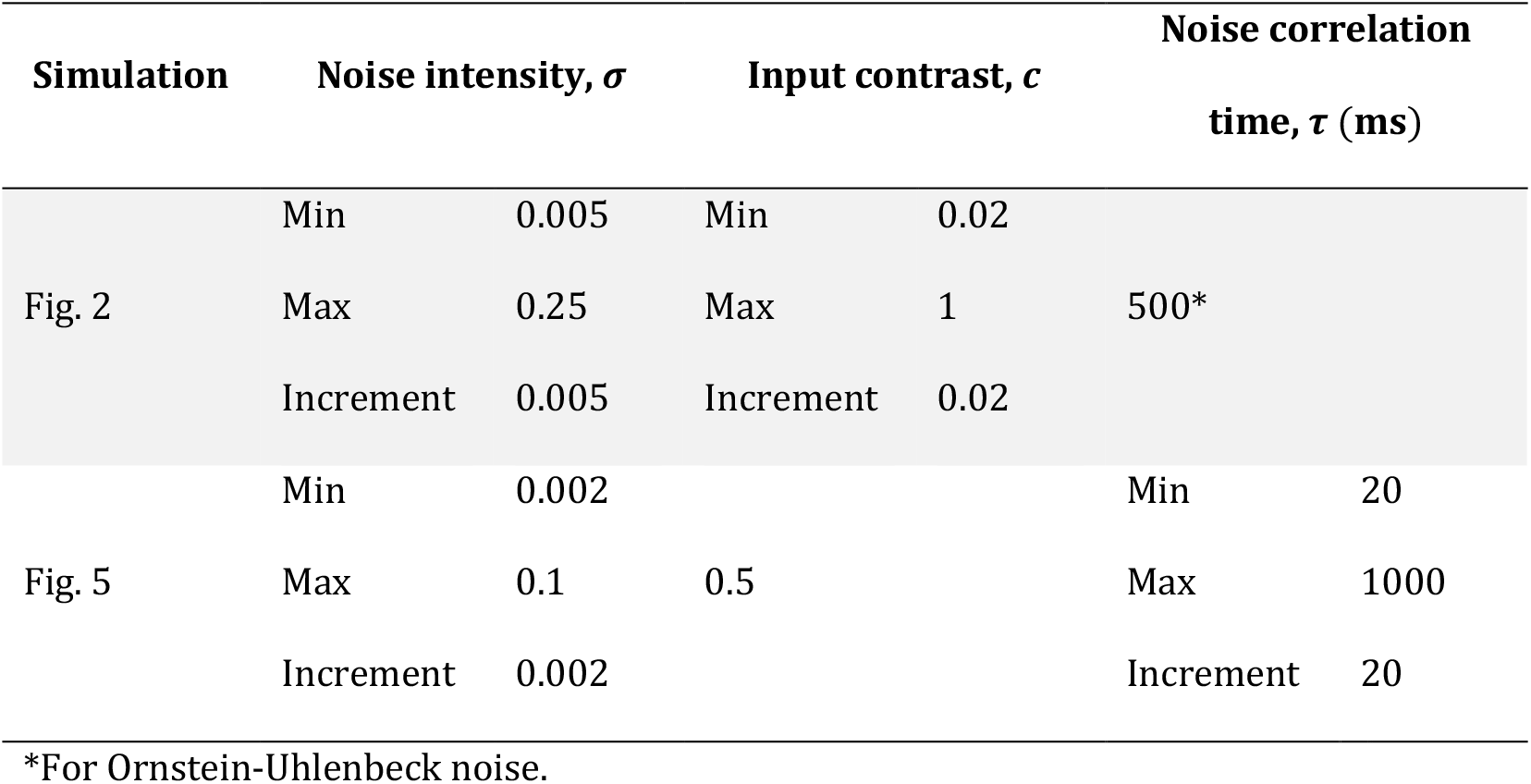
Parameter values used for the simulations in Figs. 2 and 5.

**Table 3:** Parameter values that were kept constant throughout all simulations – the same as in Said and Heeger (33).

| Parameter | Value |
| --- | --- |
| Semi-saturation constant for opponency neurons*, $s_{opp}$ | 0.9 |
| Semi-saturation constant for other neurons*, $s$ | 0.5 |
| Time constant of synaptic process, $\tau_s$ | 50 ms |
| Time constant of firing process, $\tau_r$ | 50 ms |
| Weights of network connections, $w_b$ | 1 |

\*In the normalization equation, Eq. 3.

## Supporting information

**S1 Fig. Only one formulation of adaptation results in strong dominance for white noise**. Effect of noise intensity and input contrast on relative dominance time, for simulations with Gaussian white noise and six different mathematical implementations of adaptation. Formulation 1 corresponds to the equations of divisive adaptation used in Fig. 2 and formulation 2 corresponds to subtractive adaptation. Formulations 3-6 correspond to slight variations of divisive normalization (see Methods). In all simulations, adaptation parameters were set to *w*_*H*_ = 2 and *τ*_*H*_ = 2000 ms.

**S2 Fig. Different combinations of divisive adaptation parameters do not lead to strong dominance**. Effect of noise intensity, input contrast, adaptation time constant and adaptation weight on relative dominance time, for simulations with Gaussian white noise and divisive adaptation. For each combination of noise intensity and input contrast, a diagram like the one on the right was calculated by varying the adaptation time constant and the adaptation weight. No combination of parameters gave rise to RDT above 29%. The red square on the right marks the parameter values used in Fig. S1.

**S1 Table. Simulated binocular rivalry metrics for all 9 versions of the model**. Binocular rivalry metrics for simulations with different noise processes without adaptation, with divisive adaptation and with subtractive adaptation (see Fig. 3). Only cases with RDT > 50% and for which each metric is defined (see Methods) are included, resulting in different N values. For Gaussian white noise without adaptation and with divisive adaptation, there are no cases with RDT > 50%, so this constraint was lifted and RDT > 20% was used instead.

